# Characteristics of intestinal microflora and dysbiosis in relation to the disease duration in patients with Meniere’s disease

**DOI:** 10.1101/2022.05.06.490877

**Authors:** Fumihiro Mochizuki, Manabu Komori, Yoshiyuki Sasano, Yusuke Ito, Michael E Hoffer, Izumi Koizuka

## Abstract

**Hypothesis:** Dysbiosis of the intestinal microflora has been reported in stress-induced depression and irritable colitis, a similar abnormality may also occur in stress-triggered Meniere’s disease.

**Background:** Meniere’s disease is an intractable disease characterized by paroxysms of intense rotatory dizziness, hearing loss, and other auditory symptoms. It is believed to be caused by endolymphatic hydrops of the vestibule and cochlea. The diagnosis of Meniere’s disease is based on the diagnostic criteria established by the Barany Society and AAO-HNS and Journal of Vestibular Research as definite or probable cases. Endolymphatic hydrops identification using cochlear contrast-enhanced MRI (hybrid of reversed image of positive endolymph signal and native image of positive perilymph signal [HYDROPS]) is an objective test for Meniere’s disease, enabling reliable diagnosis of this disease.

**Methods:** We investigated the gut microbiota of 10 patients (6 males and 4 females, mean age 49.6 ± 8.1 years) who met the diagnostic criteria for a unilateral definite case of Meniere’s disease using the diagnostic criteria of the Journal of Vestibular Research, and who also had significant endolymphatic hydrops on the affected side on HYDROPS. Intestinal microbiota tests were performed on these 10 patients and the results were evaluated in relation to the duration of disease, results of audiometry of the affected side, and DHI (Dizziness Handicap Inventory) score.

**Results:** A significant negative correlation was found between the disease duration and Shannon diversity index and Faith’s phylogenetic diversity, which indicated dysbiosis of the intestinal microflora. No correlation was found between the indicators of microbial diversity and the results of audiometry or the DHI on the affected side. Dysbiosis of the intestinal microbiota worsened with increasing duration of Meniere’s disease. Moreover, *Akkermansia muciniphila* was not detected in any patient with Meniere’s disease.

**Conclusions.:** Despite the small number of cases in this study (n = 10), the findings indicate the possibility of abnormalities of the intestinal microflora in Meniere’s disease.

## Introduction

Meniere’s disease is an intractable disorder characterized by paroxysms of intense rotatory vertigo and auditory symptoms such as aural fullness and hearing loss. Although the pathogenesis of Meniere’s disease is not fully understood, it is speculated to be caused by endolymphatic hydrops of the vestibule and cochlea. The diagnostic criteria for Meniere’s disease were defined by the Barany Society and AAO-HNS and Journal of Vestibular Research,etc [1]. Objective tests for evaluation of Meniere’s disease traditionally included the furosemide test and cochleography, but imaging studies using cochlear contrast-enhanced MRI (hybrid of reversed image of positive endolymph signal and native image of positive perilymph signal [HYDROPS]) have made it possible to visually evaluate endolymphatic hydrops.

HYDROPS revealed that the presence of endolymphatic hydrops on imaging is not necessarily consistent with the pathophysiology of Meniere’s disease. Appearance of could be observed in Meniere’s disease patients both in symptomatic and asymptomatic ears. [2,3].Meniere’s disease is considered a result of multiple genes interacting with environmental factors.[2,4] Several abnormalities have been reported as histological features of Meniere’s disease. These include findings of ischemia of the stria vascularis, fibrous tissue proliferation in the saccular, atrophy of the sac and loss of epithelial integrity, hypoplasia of the vestibular aqueduct, and spiral ganglion degeneration at the apex of the cochlea.[2,5] In addition, fluid homeostasis in the inner ear, as in the kidney, depends on water production, transport, and absorption. Therefore, arginine-vasopressin (AVP) localized in the inner ear, AVP-related molecules such as vasopressin receptors, and aquaporin (AQP), a water channel, are associated with its homeostasis. In animal,the action of AVP on inner ear tissues caused endolymphatic edema of the inner ear [6,7,8]. In other words, these abnormalities in inner ear homeostasis may be involved in causing endolymphatic hydrops. This AVP is known to be elevated by stressors, but there is controversy. [9] A study of patients with refractory Meniere’s disease, in which plasma AVP was measured before and after surgery, reported that plasma AVP was lower after surgery than before surgery, indicating better control of Meniere’s disease. [6] This author states that avoiding stress as much as possible is important in controlling Meniere’s disease.

Treatment of Meniere’s disease consists of daily lifestyle guidance and medication for mild cases and surgical approaches, such as intratympanic corticosteroids or intratympanic gentamicin or endolymphatic sac release or destructive surgery, for moderate to severe cases.[2]

DNA testing with next-generation sequencing has made it possible to investigate the intestinal microbiota in fecal material in great detail. As a result, the changes in the intestinal microbiota have been tested in relation to various disease groups. In this regard, various studies have investigated the intestinal microbiota of patients with depression and irritable bowel syndrome, in which stress factors are associated with the induction of pathological conditions. Ménière’s disease is a disease that repeatedly worsens and improves. In the course of treating many patients with Meniere’s disease, the authors observed several patients with complaints of diarrhea, constipation, or other abnormalities despite the absence of pre-existing bowel disease such as IBS during exacerbations. We hypothesized that the gut microbiota of Ménière’s disease patients may differ from that of healthy individuals. To test this hypothesis, we investigated the intestinal microbiota of 10 patients who met the diagnostic criteria for a solid case and had significant endolymphatic edema on the affected side by HYDROPS, an objective test.

## Materials and Methods

### Participants

Ten patients (6 males, 4 females; mean age, 49.6 ± 8.1 years) with unilateral definite Meniere’s disease diagnosed according to the diagnostic criteria of Journal of Vestibular Research [1] and showing significant endolymphatic hydrops on the affected side on HYDROPS of the inner ear were included in this study. The HYDROPS images are presented in Fig 1. The patients did not have any psychiatric, diabetic, or gastrointestinal diseases that may have caused dysbiosis.

**Fig 1.**
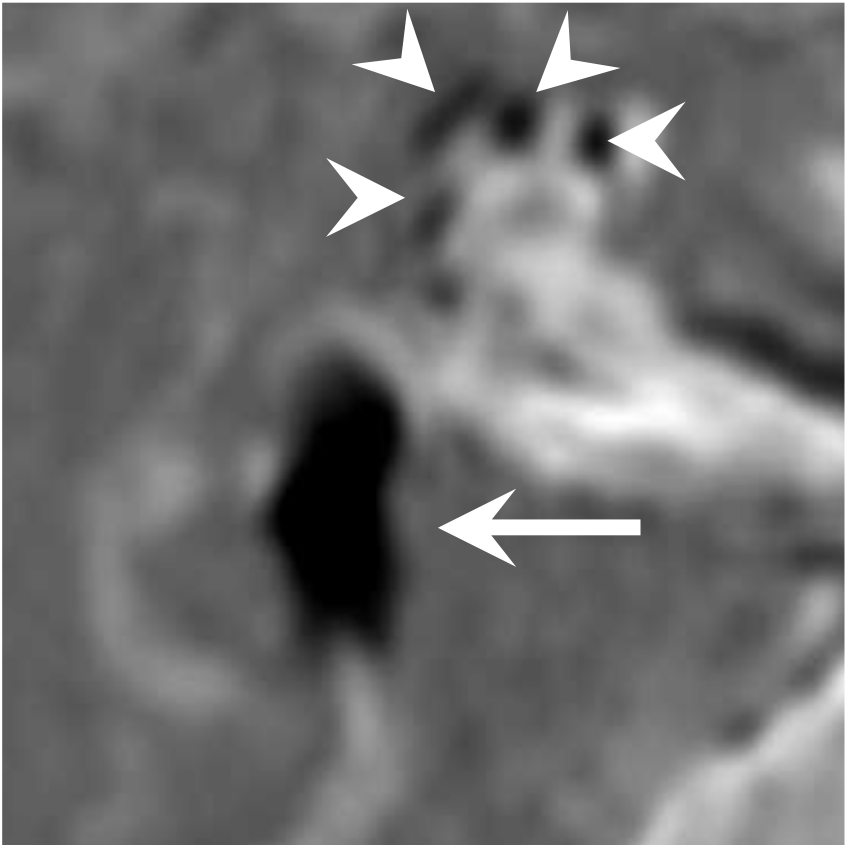
HYDROPS image in case 1. The short arrows indicate the cochlea, while the long arrow indicates the vestibule. Short arrow is cochlea, Long arrow is vestibular

## Methods

This study was approved and conducted as an observational study by the Clinical Trials Subcommittee (Observational Studies) of St. Marianna University School of Medicine. The approval number is 5674. Informed consent was given to the subjects in writing and orally. The subject’s consent was obtained orally and documented in the medical record. The information obtained was anonymized so that individual patients could not be identified.

The 10 participants received a fecal collection kit and obtained fecal samples during a period with no abnormal bowel movements. Intestinal microflora testing was performed by an external contractor (SAIKINSO Co., Tokyo, Japan). The intestinal microflora was analyzed as follows.

### Fecal sampling, DNA extraction, and sequencing

Fecal samples were collected using Cykinso fecal collection kits containing guanidine thiocyanate solution (Cykinso Inc., Tokyo. Japan) and stored at 4°C. DNA extraction from the fecal samples was performed using an automated DNA extraction machine (GENE PREP STAR PI-480; Kurabo Industries Ltd., Osaka). The V1-V2 region of the 16S rRNA gene was amplified using a forward primer (16S_27Fmod: TCG TCG GCA GCG TCA GAT GTG TAT AAG AGA CAG AGR GTT TGA TYM TGG CTC AG) and reverse primer (16S_338R: GTC TCG TGG GCT CGG AGA TGT GTA TAA GAG ACA GTG CTG CCT CCC GTA GGA GT) with the KAPA HiFi Hot Start Ready Mix (Roche). To sequence the 16S amplicons using the Illumina MiSeq platform, dual index adapters were attached using the Nextera XT Index kit. The DNA concentration of the mixed libraries was quantified by qPCR using KAPA SYBR FAST qPCR. The master mix (KK4601, KAPA Biosystems) was used with primer 1 (AAT GAT ACG GCG ACC ACC) and primer 2 (CAA GCA GAA GAC GGC ATA CGA). Library preparations were performed according to the Illumina 16S library preparation protocol (Illumina, San Diego, CA, USA). Libraries were sequenced using the MiSeq Reagent Kit v2 (500 cycles) and 250-bp paired ends.

### Taxonomy assignment based on 16S rRNA gene sequences

The paired-end reads of the partial 16S rRNA gene sequences were analyzed using QIIME 2 (version 2020.8). The steps for data processing and assignment based on the QIIME 2 pipeline were as follows: (1) DADA2 was used for joining paired-end reads, filtering, and denoising; (2) taxonomic information was assigned to each ASV by using a naive Bayes classifier in the QIIME 2 classifier with the 16S gene of the V1-V2 region data of SILVA (version 138) to determine the identity and composition of the bacterial genera. The reliability of these tests has been reported previously [10,11].

Shannon diversity index (Shannon index) and Faith’s phylogenetic diversity (FPD) were used as indicators of dysbiosis in the gut microbiota. These indices were averaged over 10000 reads per sample. The Dizziness Handicap Inventory (DHI) is a standard questionnaire that quantitatively evaluates the degree of handicap in the daily life of patients with vestibular disorders; it consists of 25 questions [12,13]. The total score ranges from 0 (no disability) to 100 (severe disability). Hearing test is that pure tone audiogram thresholds were measured at octave intervals between 125 Hz and 8 kHz. The results of the audiometric tests that were compared used a quadrant method in which the hearing levels at frequencies of 500, 1,000, and 2,000 Hz were a, b, and c dB, respectively, and the value (dB) was calculated using the formula (a+2b+c)/4.

We performed statistical analysis of the obtained gut microbiota data, age, BMI, duration of Meniere’s disease, DHI results, and hearing test results (quadrant method) by using Spearman’s correlation coefficient values. The hearing test and DHI evaluation performed closest to the intestinal microbiota tests were used for analyses. Excel with add-in software Stacel4 (OMS, Tokyo, Japan) was used as statistical software.

## Results

Table 1 shows the age, sex, BMI, duration of Meniere’s disease, DHI results, hearing test results, and dysbiosis index of intestinal bacteria. A significant negative correlation was observed between disease duration and the Shannon index (*r* = -0.72, *p* = 0.02) and FPD (*r* = -0.89, *p* = 0.0005) (Figs 2 and 3). However, no significant correlation was found between age and Shannon index (*r* = 0.16, *p* = 0.64) or FPD (*r* = -0.35, *p* = 0.31), BMI and the Shannon index (*r* = -0.21, *p* = 0.55) and FPD (*r* = -0.14, *p* = 0.68), the DHI results and the Shannon index (*r* = -0.42, *p* = 0.22) and FPD (*r* = -0.08, *p* = 0.81), and between the hearing test results and the Shannon index (*r* = -0.03, *p* = 0.94) and FPD (*r* = -0.28, *p* = 0.42).

**Table 1.** Detailed data on Meniere’s patients.

| **No** | **Age** | **BMI** | **Sex** | **Disease**<br>**period**<br>**(year)** | **DHI** | **Hearing**<br>**test**<br>**(dB)** | **Shannon's**<br>**index** | **FPD** |
| --- | --- | --- | --- | --- | --- | --- | --- | --- |
| **1** | 54 | 22.31 | Male | 3 | 46 | 7.5 | 6.34 | 18.41 |
| **2** | 39 | 26.78 | Male | 13 | 56 | 26.3 | 5.34 | 11.17 |
| **3** | 53 | 26.37 | Female | 0.67 | 16 | 31.3 | 6.58 | 19.69 |
| **4** | 60 | 19.81 | Female | 6 | 20 | 58.8 | 6.25 | 14.48 |
| **5** | 54 | 35.11 | Male | 7 | 8 | 26.3 | 6.02 | 15.83 |
| **6** | 47 | 25.6 | Male | 5 | 20 | 13.8 | 5.9 | 13.55 |
| **7** | 36 | 23.38 | Female | 10 | 54 | 36.3 | 5.77 | 12.48 |
| **8** | 61 | 25.65 | Female | 6 | 86 | 22.5 | 5.14 | 14.45 |
| **9** | 46 | 20.28 | Male | 0.58 | 62 | 21.3 | 6.7 | 21.12 |
| **10** | 46 | 22.91 | Male | 6 | 38 | 35 | 5.3 | 12.88 |
| **Average** | 49.6 | 24.82 |  | 5.725 | 40.6 | 27.91 | 5.934 | 15.406 |

**Fig 2.**
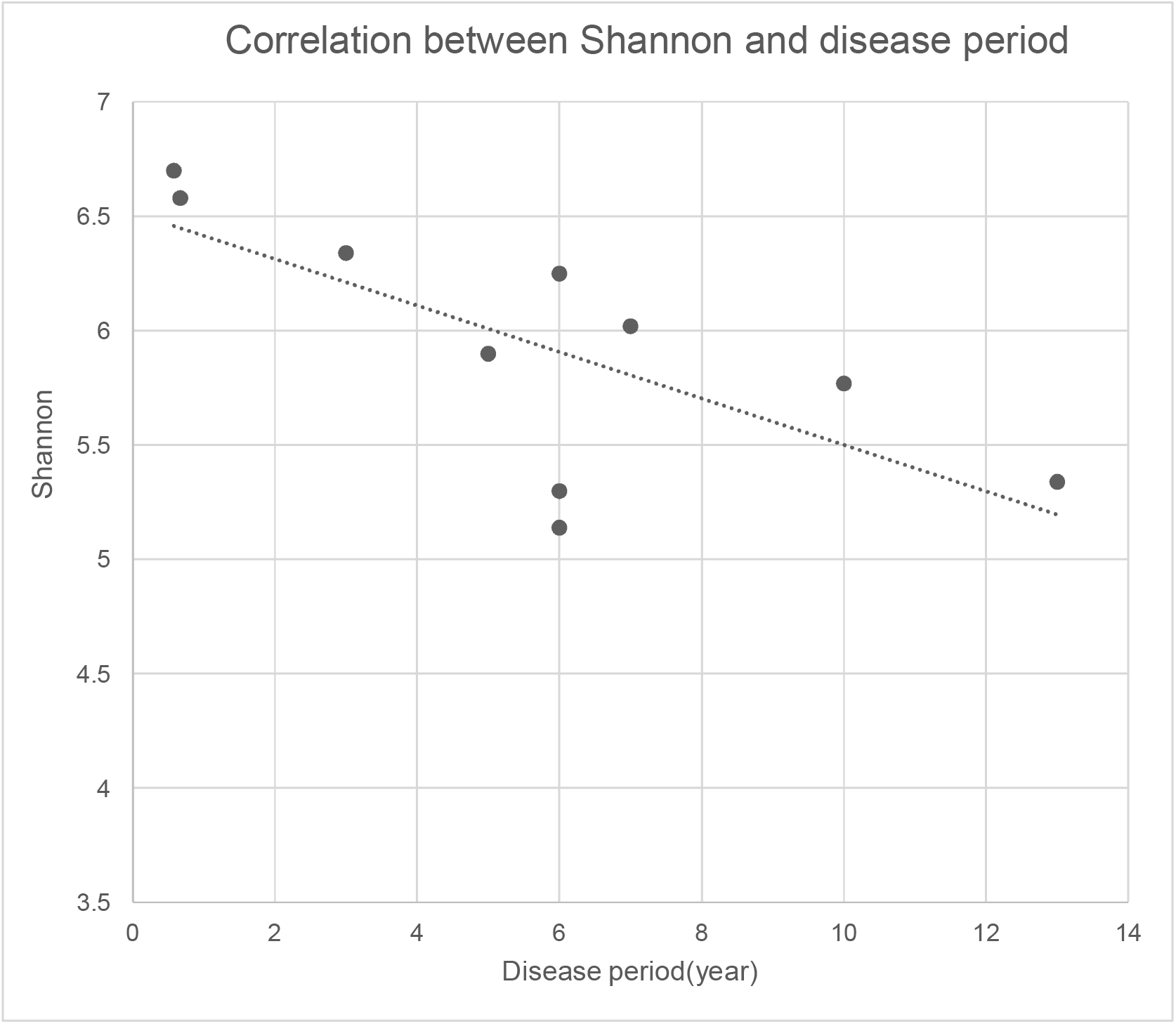
r = -0.72, p = 0.02

**Fig 3.**
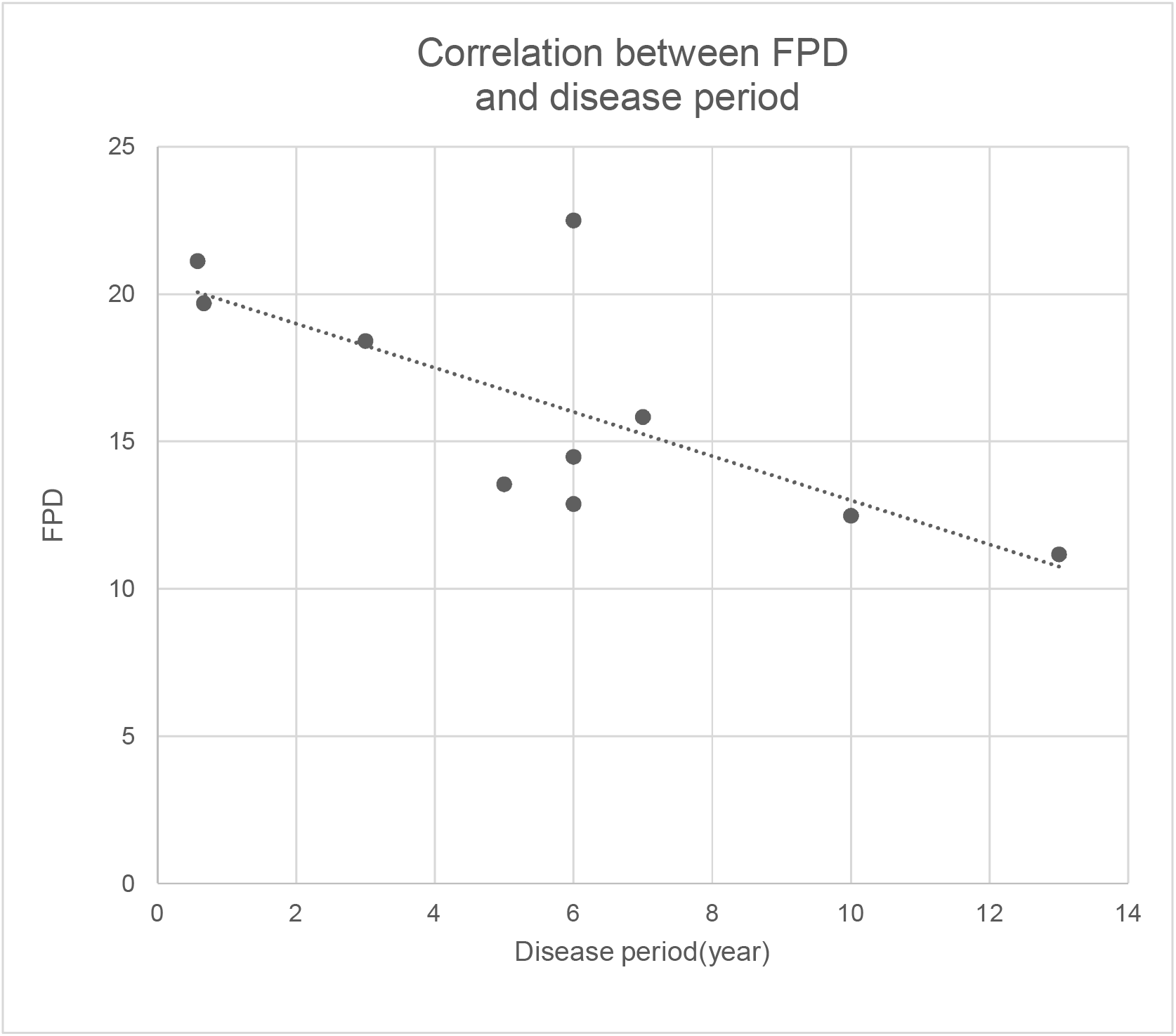
r = -0.89, p = 0.0005

The TAXON BAR showing the intestinal bacterial layer in each case is shown in Figs 4 and 5. The organisms identified by phylum, class, order, family, genus, and species are shown in Fig 4 (the description by bacterial species is too voluminous to be included), and those identified by phylum, class, and order are shown in Fig 5. *Akkermansia muciniphila* was not detected in the gut microbiota of any of the patients.

**Fig 4.**
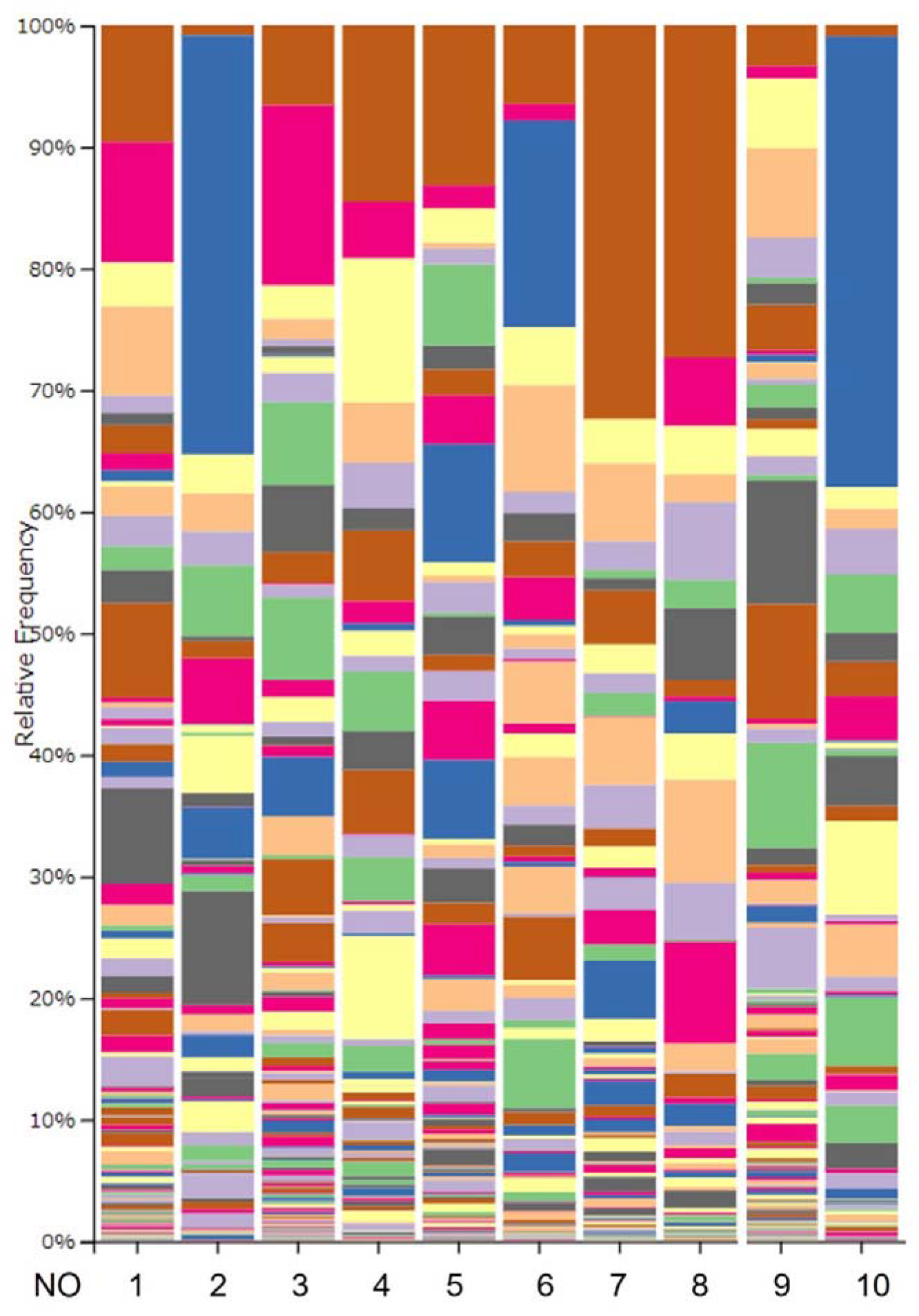
Identification by phylum, class, order, family, genus, and species.

**Fig 5.**
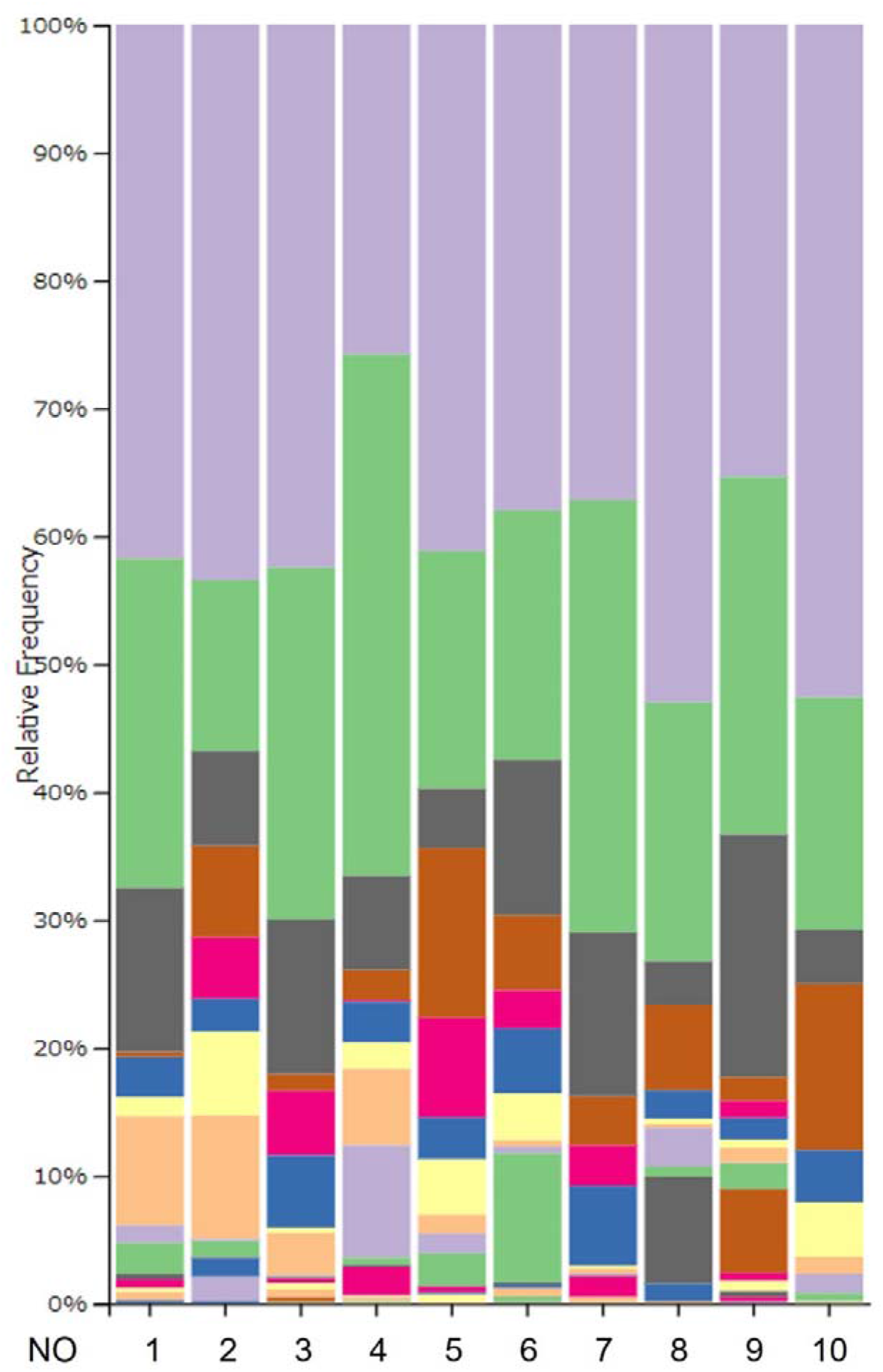
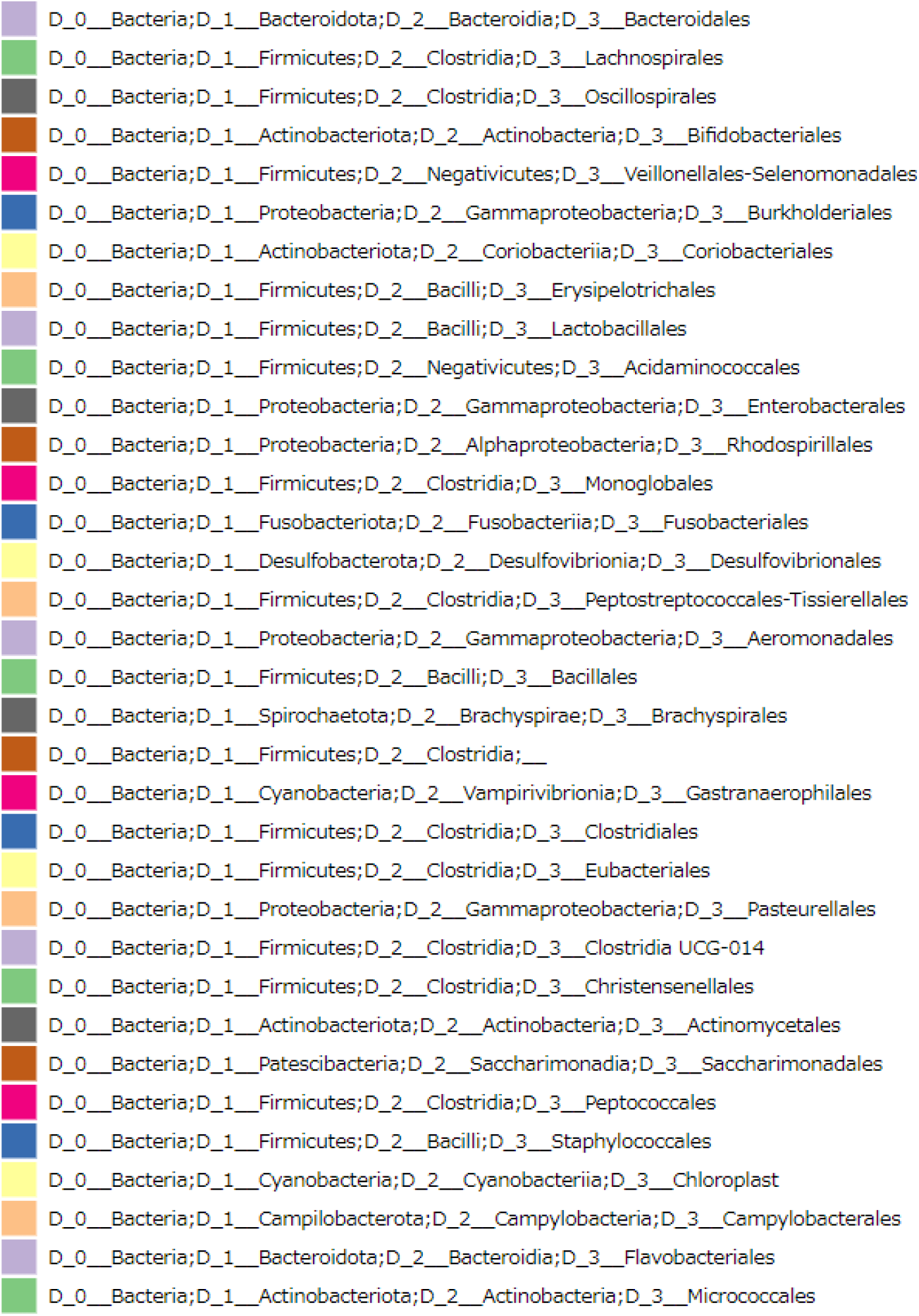
Identification by phylum, class, and order.

## Discussion

The use of gut microbiota testing to analyze various disease groups is a fairly recent approach. Recognition of the interrelationship between the brain and intestines via intestinal bacteria, which constitute the so-called microbiome-gut-brain axis, and the development of analytical scientific methods such as metagenomic analysis using next-generation sequencers and metabolome analysis have together promoted rapid accumulation of knowledge about the mechanisms underlying the microbiome-gut-brain axis.[14]

In particular, the intestinal microbiota have been extensively investigated in patients with irritable bowel syndrome (IBS) [15], and dysbiosis has also been reported to be induced in the gut microbiota of depressed patients, showing a pattern similar to that of IBS [16]. These disease groups show dysbiosis of the intestinal microbiota in comparison with healthy individuals [17]. In animal studies, when the gut microbiota of patients with depression was transplanted into gut microbiota-depleted mice, the transplanted mice exhibited characteristics of depression, both at the behavioral and physiological levels. Thus, dysbiosis of the gut microbiota was considered as a factor in the pathogenesis of depression [18]. Stressors have some influence on the pathogenesis of Meniere’s disease, as reported in a study of hormones in Meniere’s disease.[6] In fact, multiple reports have described the effects of stress and depression on Meniere’s disease. Patients with refractory Meniere’s disease are prone to depressive symptoms, with 40% reported to show neurotic symptoms and 60% showing depressive symptoms [19]. Adrenocorticotropic hormone and blood cortisol levels have been shown to be elevated due to stress in patients with Meniere’s disease, and the levels of oxytocin and vasopressin, which are known as stress hormones, are also elevated in patients with Meniere’s disease [20,21].

This study showed that dysbiosis of the intestinal microbiota progresses as the duration of Meniere’s disease increases. As mentioned above, dysbiosis of the intestinal microbiota may have progressed with increasing exposure to stress, the etiological factor, as the duration of illness increased. In this study, patients with a history of psychiatric disorders or complications were excluded. In the present study, the longer the duration of Meniere’s disease, the stronger the dysbiosis, suggesting that a prolonged Meniere’s disease status may have caused dysbiosis. But, in this study, no depression scoring was conducted, and it is unclear whether this dysbiosis was caused by the presence of depressive tendencies in patients with Meniere’s disease or whether the pathophysiology of Meniere’s disease itself is a factor causing the dysbiosis. In order to clarify these issues, it is advisable to test the same patients at different times and compare the results with the depression scoring. The microbial diversity indices showed no correlation with DHI and hearing test results. Meniere’s disease is known to cause exacerbation and amelioration, but there may have been no association between the degree of disability due to Meniere’s disease and dysbiosis of the gut microbiota. However, this study was conducted on a limited number of cases, and future comparisons based on multiple examinations in the same patients and statistical analyses with a larger number of cases are necessary in this regard. We could not find any reports on the association between Meniere’s disease and intestinal microflora, or between Meniere’s disease and IBS.

Dysbiosis of the intestinal microbiota has been verified to occur first or at the onset of various diseases. For example, rheumatoid arthritis is triggered by an immune response to specific intestinal bacteria [22]. Future studies should aim to evaluate a larger number of cases and investigate whether a specific intestinal bacterium is a trigger for Meniere’s disease.

Notably, none of the patients with Meniere’s disease had *Akkermansia muciniphila,* a gram-negative enterobacterium belonging to the phylum *Verrucomicrobia* that has been identified in the feces of healthy individuals. *Akkermansia muciniphila* is capable of growing on mucin covering the gastrointestinal mucosa as a single source of nutrition [23]. It is a commensal bacterium characterized by its degradative ability and accounts for 1%-4% of the human intestinal microflora [24]. It has been suggested that *akkermansia muciniphila* is associated with metabolic syndrome such as obesity and diabetes, and *akkermansia muciniphila* is of interest as a probiotic that can ameliorate metabolic syndrome. In relation to obesity, the intestinal mucus layer of mice fed a high-fat diet that developed metabolic syndrome became thin and *akkermansia muciniphila* decreased, but oral administration of *akkermansia muciniphila* restored the thickness and permeability of the mucus layer in the intestinal tract and also improved metabolic syndrome The metabolic syndrome was reported to be improved by oral administration of *akkermansia muciniphila*[25]. In addition, human observational studies have reported that *akkermansia muciniphila* is reduced in patients with type II diabetes and atopic dermatitis compared to healthy controls[26,27]. *Akkermansia muciniphila* has been shown to improve insulin resistance and have immunomodulatory effects[25].

In mice, walking exercise was reported to increase *Akkermansia muciniphila* in the intestinal microflora. [28] In humans, increased *Akkermansia muciniphila* of the intestinal microflora has been reported with exercise and dietary restriction. [29] A study in mice reported an increase in *Akkermansia muciniphila* and *Lactobacillus* when melatonin was administered to improve sleep deprivation[30]. These reports indicate that *Akkermansia muciniphila* increases or decreases with exercise, diet, and sleep in the intestinal microflora. Daily lifestyle guidance for Meniere’s disease generally includes improving sleep, diet, and avoidance of stress [31]. Aerobic exercise is also reported to be effective for Meniere’s disease [32].

These lifestyle guidance may be consistent with some of the factors that increase *Akkermansia muciniphila* in the intestinal microflora. In the future, testing of the intestinal microflora before and after daily lifestyle guidance in patients with Meniere’s disease will clarify the relationship between *Akkermansia muciniphila* and Meniere’s disease.

Only 10 patients were evaluated in this study, and future studies should aim to accumulate data from more cases. However, the fact that dysbiosis of the intestinal microbiota depended on the duration of Meniere’s disease and that *Akkermansia muciniphila* was not detected in all cases suggests that the intestinal microbiota in patients with Meniere’s disease is different from that in normal individuals. Future investigation of the intestinal microflora in Meniere’s disease may lead to elucidation of the pathogenesis and development of new treatments. Further studies will be required to accumulate data from more cases.

## Conclusions

Ten patients with a definite diagnosis of Meniere’s disease and marked endolymphatic hydrops on HYDROPS underwent examinations of the intestinal microflora. A significant negative correlation was found between the duration of Meniere’s disease and dysbiosis of intestinal microbiota. None of the patients harbored *Akkermansia muciniphila.* However, this study was limited by the small number of patients. Nevertheless, the relationship between Meniere’s disease and the intestinal microbiota has not been reported previously, and these findings may help researchers determine the pathogenesis of Meniere’s disease and propose new treatments for it in the future.

The funders had no role in study design, data collection and analysis, decision to publish, or preparation of the manuscript.

